# Subsurface Hydrocarbon Degradation Strategies in Low- and High-Sulfate Coal Seam Communities Identified with Activity-Based Metagenomics

**DOI:** 10.1101/2021.08.26.457739

**Authors:** Hannah Schweitzer, Heidi Smith, Elliott P. Barnhart, Luke McKay, Robin Gerlach, Alfred B. Cunningham, Rex R. Malmstrom, Danielle Goudeau, Matthew W. Fields

**Affiliations:** Center for Biofilm Engineering, Montana State University, Bozeman, MT 59717, USA; Department of Microbiology & Immunology, Montana State University, Bozeman, MT 59717, USA; US Geological Survey, Wyoming-Montana Water Science Center, Helena, MT 59601,USA; Department of Land Resources and Environmental Sciences, Montana State University, Bozeman, MT 59717, USA; Energy Research Institute, Montana State University, Bozeman, MT 59717, USA; Department of Biological and Chemical Engineering, Montana State University, Bozeman, MT 59717, USA; Department of Civil Engineering, Montana State University, Bozeman, MT 59717, USA; DOE Joint Genome Institute, Berkeley, CA 94720, USA

## Abstract

Environmentally relevant metagenomes and BONCAT-FACS derived translationally active metagenomes from Powder River Basin coal seams were investigated to elucidate potential genes and functional groups involved in hydrocarbon degradation to methane in coal seams with high- and low-sulfate levels. An advanced subsurface environmental sampler allowed the establishment of coal-associated microbial communities under *in situ* conditions for metagenomic analyses from environmental and translationally active populations. Metagenomic sequencing demonstrated that biosurfactants, aerobic dioxygenases, and anaerobic phenol degradation pathways were present in active populations across the sampled coal seams. In particular, results suggested the importance of anaerobic degradation pathways under high-sulfate conditions with an emphasis on fumarate addition. Under low-sulfate conditions, a mixture of both aerobic and anaerobic pathways were observed but with a predominance of aerobic dioxygenases. The putative low-molecular weight biosurfactant, lichysein, appeared to play a more important role compared to rhamnolipids. The methods used in this study—subsurface environmental samplers in combination with metagenomic sequencing of both total and translationally active metagenomes—offer a deeper and environmentally relevant perspective on community genetic potential from coal seams poised at different redox conditions broadening the understanding of degradation strategies for subsurface carbon.

**One Sentence Summary:** Identifying hydrocarbon degradation strategies across different redox conditions via metagenomic analysis of environmental and translationally active (BONCAT-FACS) samples from subsurface coal beds.

## Introduction

The terrestrial subsurface contains the majority of Earth’s organic carbon (∼90%)^1^, and much of the carbon can be converted to methane under anaerobic conditions through biogasification (*i*.*e*., biological decomposition of organic matter into methane and secondary gases). Biogasification can take place in coal, black shale, and petroleum reservoirs and is estimated to account for over 20% of the world’s natural gas resources^2^. Factors influencing biogasification include coal rank, redox conditions (*e*.*g*., sulfate and oxygen), and the genetic potential and activity of the microbial community. Coal is a heterogeneous and highly complex hydrocarbon consisting of polycyclic aromatic hydrocarbons, alkylated benzenes, and long and short chain n-alkanes^3^, and despite the recalcitrant nature of coal, degradation by microbial consortia has been shown in a variety of coal formations^4–6^. It is generally accepted that shallower coal beds that contain sulfate do not produce methane because sulfate-reducing bacteria (SRB) outcompete methanogens for substrates (*e*.*g*., acetate, CO_2_ and hydrogen)^5,7^. In methanogenic coal beds^5,8^, hydrogenotrophic and acetoclastic methanogens are commonly identified, including different types of acetoclastic methanogens (*e*.*g*., *Methanothrix, Methanosarcina*), which have distinct pathways for acetate utilization. It remains unknown what type of methanogenesis predominates *in situ* for different coal seams under different physicochemical conditions^9,10^.

New coal degradation pathways are still being discovered and the involvement of different pathways in the turnover of refractory carbon under various redox conditions (*e*.*g*., sulfate) remains largely unresolved^11–13^. The majority of coal degradation research has focused on fumarate addition, while less is known about alternate coal degradation strategies such as phenol degradation by carboxylation and hydroxylation of alkanes, benzene and ethylbenzene^14,15^. The fumarate addition pathway involves the activation of *n*-alkanes by the addition of fumarate via the double bonds at the terminal or subterminal carbon^14,16–20^. Several fumarate addition genes (*e*.*g*., *ass*-alkylsuccinate synthase for alkanes, *bss*-benzylsuccinate synthase for alkylbenzenes, and *nms*-naphylmethylsuccinate synthase) are often used as catabolic biomarkers for anaerobic hydrocarbon degradation^16–18^. These genes have been characterized from many subsurface hydrocarbon-containing environments,^14,17,21–26^ but the importance under different redox conditions is still unclear. Carboxylation and hydroxylation strategies are less well documented mechanisms of anaerobic degradation, although, in recent years work has begun to suggest importance in anaerobic hydrocarbon degradation^15,19,27,28^, yet how these strategies vary across redox conditions *in situ* and detecting organisms responsible for degradation warrants further investigation.

While biosurfactants have not been identified *in situ* in coal seams and are not considered a necessary hydrocarbon degradation gene, previous laboratory-based research demonstrates a potentially important role in decreasing the hydrophobicity of the solid coal surface, allowing for cellular and/or protein interactions at the coal surface^29^. Therefore, biosurfactant-producing microorganisms likely play direct and indirect roles in hydrocarbon degradation^29–32^. The accumulation of the esterase hydrolase enzyme has been correlated to biosurfactant production and is often used as a biomarker for biosurfactant production^33^. Biosurfactants are routinely observed in environments that consist of complex hydrocarbons and are therefore hypothesized to be an interdependent, complex, and coordinated means of increasing coal bioavailability. However, studies that have demonstrated active biosurfactant-producing microorganisms in coal environments are lacking.

Aerobic hydrocarbon degradation is the most documented form of hydrocarbon degradation via the aerobic activation of alkanes with dioxygenase enzymes that use oxygen as an electron acceptor and as a reactant in hydroxylation^34^. Aerobic hydrocarbon degradation in coal environments is often disputed due to the uncertainty of the presence of oxygen and more research into anaerobic hydrocarbon degradation strategies has been performed in the last few decades. However, recent metagenomic analyses have indicated the presence of aerobic hydrocarbon degradation genes in coal beds across Canada, but it is unknown whether these genes were present in active organisms *in situ*^24^.

The rate-limiting step in coal biogasification has been attributed to initial biological breakdown of the refractory hydrocarbon matrix^35^. As discussed above, while several degradation strategies including the role of biosurfactants and aerobic hydrocarbon degradation have been observed, it remains challenging to link genetic potential to activity using traditional sequencing methodologies^36^. While traditional sequencing methodologies allow for an environmental snapshot of detectable genes, dead, dormant, or active groups are difficult to delineate.

In the described study, environmentally relevant samples from three separate coal seams located in the Powder River Basin (PRB) near Birney, Montana, were obtained with a Subsurface Environmental Sampler (SES; Patent # US10704993B2)^37^. The SES is a advanced sampling tool which enabled harvesting of *in situ* coal-associated biofilms representative of local environmental conditions. The SES-retrieved samples were analyzed via total and activity-based metagenomes (using BONCAT-Bioorthogonal noncanonical amino acid tagging) to examine metabolic potential across hydrologically isolated coal seams with varying redox conditions^5,8^. BONCAT methods have previously been used in combination with fluorescence-activated cell sorting (BONCAT-FACS) and SSU rRNA gene sequencing^38,39^, but metagenomic sequencing of active cells from BONCAT-FACS has yet to be performed. Total hydrocarbon degradation/surfactant potential and differences in microbial diversity from BONCAT-FACS- and shotgun-metagenomes were compared across environments with different sulfate conditions. Investigation of the active microbial groups and functional potential in coalbed methane (CBM) habitats provided new insight into the hydrocarbon degradation strategies and the associated capacities for carbon cycling that vary with depth and sulfate conditions through shallow coal seams.

## Results and Discussion

### Microbial communities and diversity under high and low-sulfate conditions

An SES^37^ was used to collect *in situ* coal-associated microbial communities for shotgun metagenomic sequencing in three geochemically diverse coal seams (Fig. S1). Samples from a high-sulfate (Nance, N-H, 2002.4-2501.2 mg/L) coal seam and two low-sulfate (Flowers Goodale, FG-L; Terret, T-L, 0.7-16.3 mg/L) coal seams were co-assembled and used to determine differences in hydrocarbon degradation genes from the entire community and the taxonomic diversity was accessed with the high quality (>80%) metagenome-assembled genomes (MAGs) (Fig. 1). There were 769,799 genes contained in all of the environmental metagenomes on 92,729 contigs. After refinement, 86 MAGs had >80% estimated completeness for the co-assembled environmental shotgun metagenomes (Fig. 2, Table S1). The broad categories of gene classifications were largely similar across all samples (Supplemental Data 1).

**Figure 1.**
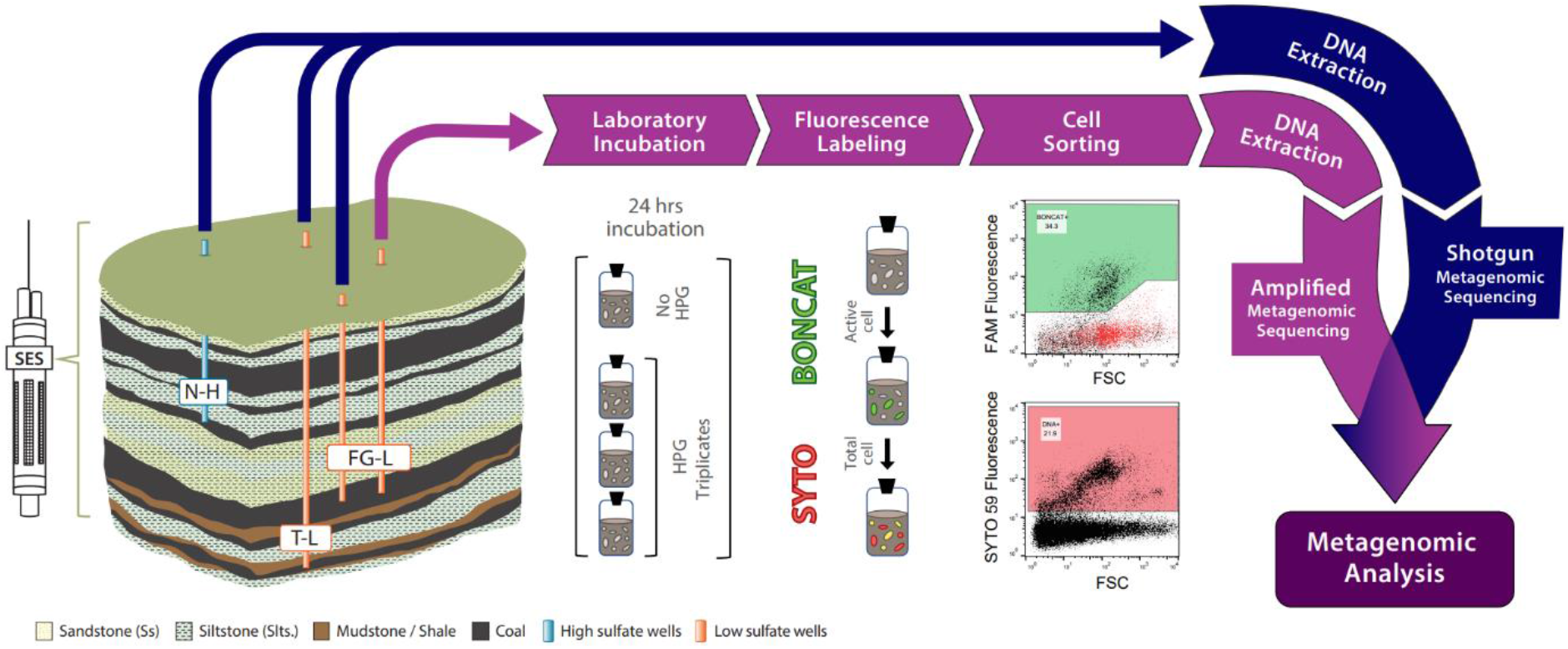
Schematic depicting experimental workflow. High-sulfate (Nance, N-L) and low-sulfate (Flowers Goodale, FG-L; Terret, T-L) CBM wells were sampled utilizing a Subsurface Environmental Sampler (SES) to retrieve in situ CBM samples of the attached fraction under environmentally relevant conditions. Following retrieval, samples were either prepared for BONCAT incubation (purple track) or shotgun metagenomics (blue track). For shotgun metagenomics, DNA was extracted and sequenced. BONCAT incubations with the synthetic amino acid tracer, HPG, lasted 24 hours followed by cell removal from coal, click chemistry, staining, and cell sorting of BONCAT+ and BONCAT-Total (SYTO stained) cell fractions, DNA extraction and amplified metagenomic sequencing. The No HPG control was used to determine sorting gates for BONCAT+ events.

**Figure 2.**
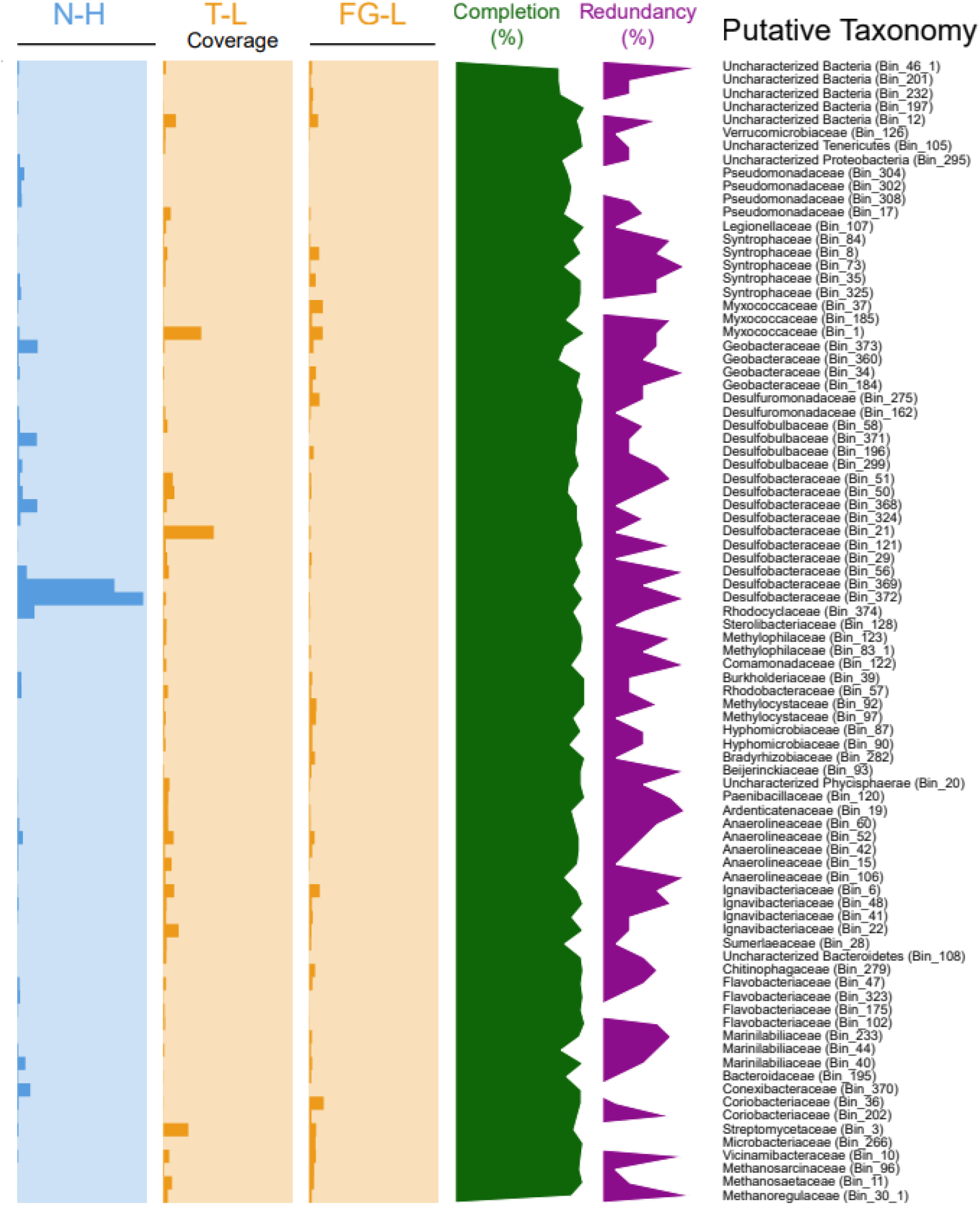
Coverage comparison of individual metagenome-assembled genomes (MAGs). MAGs with greater than 80% completion (green line graph; axis from 0-100%) from one high-sulfate coal seam (N-H in blue) and two low-sulfate coal seams (FG-L and T-L in orange) were compared. Coverage is calculated by finding the average depth of coverage across all contigs using Anvi’o compiled contigs. Coverage bar graphs axis (from 0-550X) is indicated with the highest possible sequencing depth being 550. All samples were below 10% redundancy (purple line graph; axis 0-10%). Taxonomies are listed based on phylum.

The MAGs that were most closely related to organisms from the *Desulfobacteraceae* family were observed across all sampled coal seams regardless of sulfate level, although highest coverage was in the high-sulfate coal seam (Fig. 2). The *Desulfobacteraceae* family consists of commonly studied sulfate reducing bacteria (SRB) and is known to be metabolically versatile and exhibits multiple electron transfer complexes suggesting alternative metabolic mechanisms^40^. Previous research has demonstrated increased abundance of SRBs in high-sulfate coal seams compared to low-sulfate coals seams^5^. Although, *dsr*A genes have been detected in acetate-amended methane-producing coal seams, and these results suggest organisms with *dsr*A may play an important role in syntrophic biogasification in the absence of sulfate^41^.

In the high-sulfate coal seam (N-H), the *dsr*A gene was identified in MAGs taxonomically similar to *Desulfobacteraceae* (Bin 372) and *Desulfatibacillum alkenivorans* (Bin 369), whereas the FG-L and T-L low-sulfate coals were predominated by MAGs with < 80% completion as well as *Desulfococcus (*Bin 21*), Desulfuromonas* (Bin 275), and *Myxococcaceae* (Bin 1 and Bin 37) (Fig. 3, Supplemental Data 2). Previous research on the *Desulfococcus* clade and the genus *Desulfatibacillum alkenivorans* reported the ability to anaerobically degrade alkanes^42,43^. *Desulfatibacillum alkenivorans* has also been described as being capable of using hexadecane as the sole carbon source in the presence of sulfate^43^. *Anaeromyxobacter dehalogenans* (from the *Myxococcaceae* family) is a facultative anaerobe that utilizes acetate with nitrate, oxygen, and fumarate^44^.

**Figure 3.**
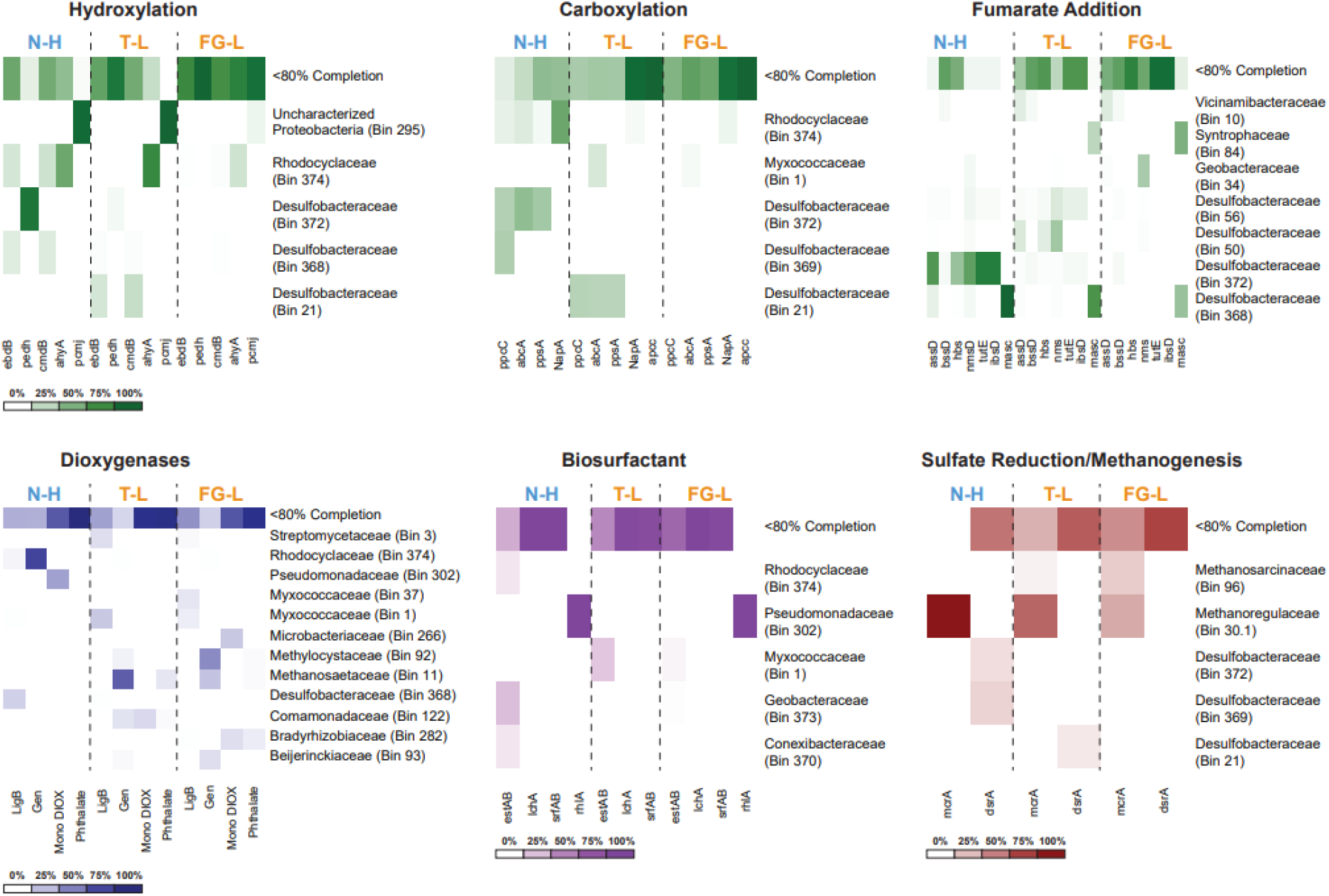
Comparison of gene abundances across metagenome assembled genomes MAGs. Heatmaps from all MAGs from shotgun metagenomic samples depicting the relative gene abundane for individual genes involved in hydrocarbon degradation for the high-sulfate coal seam (N-H) and two low-sulfate coal seams (T-L and FG-L). Only high quality MAGs (>80% completion) were taxonomically classified. All MAGS below 80% completion were grouped together as <80% Completion. The six different heatmaps, displaying the genes of interest, were grouped based on method of anaerobic degradation (green heatmaps) via hydroxylation, carboxylation, and fumarate addition, aerobic degradation (blue heatmap) via dioxygenases, biosurfactant production (purple heat map) and sulfate reduction/methanogenesis (red heatmap). All relative abundance of each gene found in the corresponding MAGs are represented by a color gradient (as seen in legend below heatmap) with the darkest color indicating the highest relative abundance for each sample and white indicating lack of the gene.

The most predominant MAGs in the low-sulfate FG-L and T-L wells were most closely related to *Anaeromyxobacter* (*Myxococcaceae*), *Desulfococcus* (*Desulfobacteraceae*), and methylotrophic organisms from the *Methylocystaceae* family consisting of *Methylocystis* and *Methylosinus* (Fig. 2 and Table S1). Methane producing organisms such as acetoclastic methanogens, *Methanothrix* (*Methanosaetaceae*) and *Methanococcoides* (*Methanosarcinaceae*), and the hydrogentorphic methanogen, *Methanoregula formicica* (*Methanoregulaceae*), were also detected in the low-sulfate coal seams (Fig. 2 and Table S1). Hydrogenotrophic and acetoclastic methanogens are commonly identified in methanogenic coal beds^5,8^, and different types of acetoclastic methanogens (*e*.*g*., *Methanothrix, Methanosarcina*) have different pathways for acetate utilization. Active acetate utilization has been demonstrated in PRB coalbeds but the dominant microbial communities or metabolic pathways that generate or utilize acetate in these environments are generally unknown^36^. As stated above, *Anaeromyxobacter dehalogenans* typically thrive in environments with acetate further supporting the possibility of acetate in the low-sulfate communities^44^. Previous laboratory investigations have demonstrated that acetoclastic methanogens produce a large fraction of biogenic methane in coal enrichments and it is therefore hypothesized that high methane producing coal seams are abundant in acetoclastic methanogens^45,46^. In comparison to other methanogens, *Methanothrix spp*. have been reported to be better scavengers in low-acetate conditions^10^. However, recent isotopic analysis of PRB samples reveal that hydrogenotrophic methanogenesis can also be a significant methane producing pathway^45^.

To further investigate the dominant methanogenic populations in the low-sulfate coal seams, the BONCAT-FACS active (BONCAT+) metagenome from the FG-L coal seam was analyzed and resulted in 24 MAGs (Fig 4, Table S2). The entire BONCAT+ library had 888 contigs with 31,748 genes, while the total SYTO stained population (BONCAT-Total) from the same experiment yielded 71,455 genes on 1,748 contigs (Fig. S2, Table S3). Comparison of the BONCAT-Total and BONCAT+ samples demonstrated that 29.3 - 36.3% of the recovered cells belonged to populations that were BONCAT+ (Fig. S3). Six of the 24 BONCAT+ MAGs were taxonomically similar to known methanogens, *Methanothrix*, an acetoclastic methanogen and *Methanobacterium*, a hydrogenotrophic methanogen (Fig. 4, Table S2). These results suggest there are likely multiple (hydrogenotrophic and acetoclastic) active methanogenic pathways *in situ*^5,45^, although the *Methanothrix* MAG exhibited significantly higher quality (Fig. 4).

**Figure 4.**
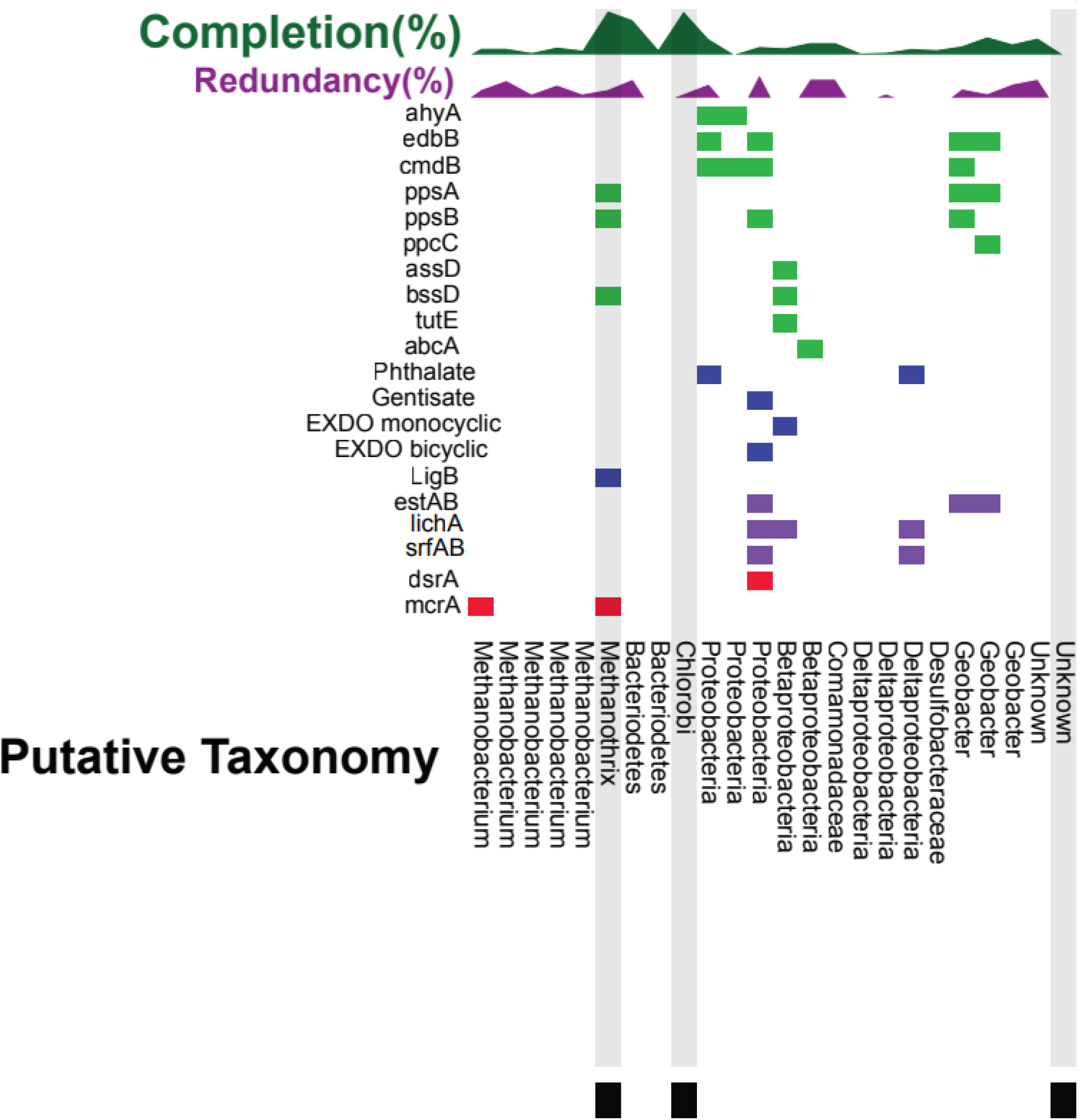
Comparison of coverage and gene presence in BONCAT positive metagenome assembled genomes (MAGs). The presence of genes for each corresponding MAG is indicated with a color coded box: anaerobic hydrocarbon degrading (green), aerobic hydrocarbon degrading (blue), biosurfactant (purple) and sulfate reduction/methanogenesis (red). The percent completion (green line graph out of 100%) and redundancy (purple line graph out of 10%) were compared for each MAG. The three highlighted MAGs with the symbol (◼) have a >99% average nucleotide identity similarity to BONCAT-Total MAGs indicated with the same symbol.

Notably, three BONCAT+ MAGs had > 70% completeness and were related to the Bacteroidetes, Chlorobi and *Methanothrix* (Fig. 4, Table S2). The Chlorobi and *Methanothrix* MAGs overlapped between the BONCAT+ and the BONCAT-Total metagenome as determined by average nucleotide identity (Fig. 4, Fig. S2, Table S2, Table S3 and Supplemental Data 3). Bacteroidetes and Chlorobi have previously been observed to dominate acetate amended coal seams, and furthermore, microorganisms belonging to the Bacteroidetes phylum have been identified as a key lineage in hydrocarbon degradation^41,47^. These high quality BONCAT+ MAGs are examined in greater metabolic detail for potential involvement in coal degradation, acetate production, and subsequent acetoclastic methanogenesis in a separate study^48^.

### Presence of Biosurfactant Genes

Presumptive biosurfactant genes for an esterase hydrolase enzyme (*estAB*), lichenysin synthetase (*lchA*), and surfactin synthetase (*srfAB*) were detected with the highest gene abundance in the FG-L coal seam (Fig. 5, Table S4). The *estAB* gene had the highest abundance (between 21.2-32.3 RPKM) compared to any of the investigated biosurfactant genes and was primarily in MAGs from the environmental metagenomes with < 80% estimated completeness (Fig. 3, Fig. 5, Table S4, and Supplemental Data 2). The *estAB* gene was also observed in three of the 24 BONCAT+ MAGs, two of which were most closely related to *Geobacter* and one related to Proteobacteria (Fig. 4, Table S2). Previous work has suggested the importance of esterases in the production of biosurfactants due to the correlation between the accumulation of the esterase hydrolase enzyme and the production of lichenysin and surfactin^33^. Lichenysin, surfactin and the esterase genes had the highest abundance in the low-sulfate coal seams and were identified in the BONCAT+ metagenomes from FG-L (Fig. 4, Fig. 5, Table S2 and Table S4).

**Figure 5.**
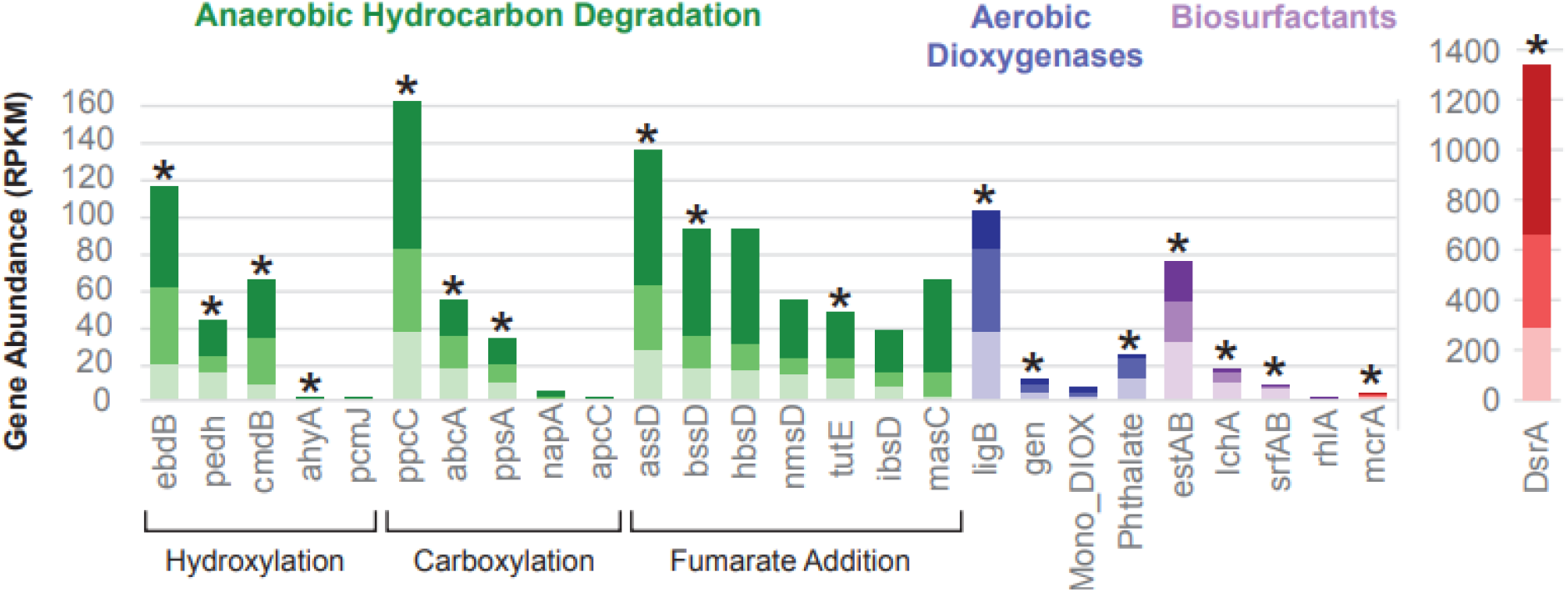
Comparison of gene abundances across coal seams. Gene abundance was calculated using the coverage of each gene per kilobase of transcript per million mapped reads (RPKM) for each gene of interest from all shotgun metagenomic samples. Representative genes for each of the databases tested are presented here, with genes of interest containing the highest gene abundance in one of the samples and/or being present in the BONCAT+ sample. The coverage of each gene was calculated using the average depth of coverage across the contig the gene was identified in. All non-duplicate contigs from the entire community containing the gene of interest above 40% identity were summed and reported here. The different genes are grouped based on anaerobic hydrocarbon degradation strategies (green), aerobic dioxygenases (blue), biosurfactant (purple) and sulfate reduction/methanogenesis genes (red). All three environmental samples are represented in a color gradient with the high sulfate coal seam, N-H in the darkest shade (top), and two low sulfate coal seams, T-L (middle) with medium shade and FG-L (bottom) in the lightest shade. Stars indicate the presence of the individual gene in the BONCAT+ sample. The DsrA gene has its own axis because it had a greater overall gene abundance.

In the low-sulfate environmental metagenomes (FG-L and T-L), the *lchA* and *srfAB* biosurfactant genes were most present in the MAGs that were < 80% estimated completeness (Fig. 3) and one highly complete (>80%) MAG that was taxonomically similar to *Legionella* (Supplemental Data 2). *Legionella* spp. have previously been shown to have the ability to produce biosurfactants that contain a lipid structure and had a similar environmental response as surfactin produced from *Bacillus subtilis*^49^. Lichenysin and surfactin are part of the surfactin operon family and contain four open reading frames that include *lchA* and *srfAB*. Much like surfactin, lichenysin is a low molecular weight anionic cyclic lipopeptide biosurfactant previously shown to be produced by *Bacillus licheniformis* isolates^50^. Previous research has also demonstrated lichenysins are capable of enhancing oil recovery and degrading naphthalene and crude oil^50,51^. Biosurfactants have been suggested as having a potentially important role in decreasing the hydrophobicity of the solid coal surface, allowing for cellular and/or protein interactions at the coal surface^29^. Both lichenysin and surfactin are nonribosomal peptides and therefore are less energy intensive and could be ideal for oligotrophic conditions such as coal seams^50,52,53^. These results suggest biosurfactants are likely important in coal biogasification in low-sulfate environments and could facilitate the initial steps in hydrocarbon breakdown.

The rhamnosyltransferase gene (*rhlA*) had the lowest gene abundance of any of the investigated biosurfactant-related genes. The greatest *rhlA* abundance was observed in the high-sulfate coal seam with little to no presence in the low-sulfate coal seams (Fig. 5, Table S4). In addition, *rhlA* was not detected in the BONCAT+ sample, although it was found in one MAG from the environmental shotgun metagenomes related to *Pseudomonadaceae* (Bin 302) (Fig. 3, Fig. 4, Fig. 5, Table S2 and Table S4). Rhamnolipid biosurfactants produced by *Pseudomonas aeruginosa* and *Pseudomonas stutzeri* have previously shown the ability to enhance anaerobic degradation of the PAHs, phenanthrene and pyrene^54–56^. *Pseudomonas stutzeri*, a model biosurfactant producing organism, was detected in formation water originating from volatile bituminous coal in the Appalachian Basin coal bed and is suggested to play an important role in enhanced oil recovery^55–57^. The greater prevalence of *lchA* and *srfAB* suggests that rhamnolipids may not be the dominant biosurfactants in all coal environments as previously hypothesized from benchtop coal enrichment experiments^54–57^.

### Hydrocarbon Degradation in High-Sulfate Environments

The phenylphosphate carboxylase gene (*ppc*C) had the highest abundance in the N-H coal seam (Fig. 5, Table S4). Phenylphosphate carboxylase and phenylphosphate synthase can contribute to the anaerobic degradation of phenol, and phenylphosphate carboxylase is responsible for the conversion of phenylphosphate and CO_2_ into 4-hydroxybenzoate and has previously been identified in *Geobacter metallireducens*^58^. Consistent with this, one of the most prominent populations represented by MAGs with > 80% estimated completeness in the high-sulfate coal seam (N-H) was taxonomically similar to *Geobacter* (Bin 360, Bin 373, Bin 34 and Bin 184) and contained the *ppc*C gene (Fig. 2, and Supplemental Data 2).

The gene with the next highest abundance for the high-sulfate coal seam was the alkylsuccinate synthase subunit D (*ass*D) followed by hydroxybenzylsuccinate synthase (*hbs*D), and benzylsuccinate synthase subunit D (*bss*D) (Fig. 5, Table S4). These genes (*ass*D, *hbs*D and *bss*D) are all involved in hydrocarbon degradation via fumarate addition, which is currently the only well-described, oxygen-independent alkane activation^59^. For the high-sulfate coal seam, fumarate addition genes were detected in MAGs that were most taxonomically similar to populations from the *Desulfobacteraceae* family (Bin 372) as well as others such as *Desulfatibacillum alkenivorans* (Bin 56) and the *Desulfococcus* clade (Bin 50, Bin 368) (Fig. 3). Previous research has shown these alkane-degrading organisms in the presence of sulfate^42,43^. Our results indicate fumarate addition may play a more crucial role in high-sulfate than low-sulfate coal seam environments for hydrocarbon degradation under high-sulfate conditions.

### Low-sulfate hydrocarbon degradation: Aerobic and Anaerobic Strategies

Previously reported methane concentrations for low-sulfate FG-L and T-L (33-67 mg/L^8^) are consistent with our detection of higher *mcr*A gene abundances compared to the high-sulfate (< 0.15 mg/L methane) coal seam (Fig. 3, Fig. 5, Table S4). The rate-limiting step in coal biogasification has been attributed to initial biological breakdown of the refractory hydrocarbon matrix^35^. Analysis of the hydrocarbon degradation genes showed a mix of both aerobic and anaerobic strategies, yet there was a higher gene abundance of aerobic hydrocarbon degradation genes under low-sulfate compared to the high-sulfate coal seam. The homoprotocatechuate dioxygenase α subunit gene (*lig*B) was present in the BONCAT+ metagenome (Fig. 4, Table S2, and Table S4), and in the environmental metagenomes. The *lig*B had the highest abundance of all investigated aerobic aromatic hydrocarbon degradation genes^60^ regardless of sulfate level. Moreover, *lig*B had the highest gene abundance for any gene detected in the low-sulfate coal seams (Fig. 5, Table S4).

Homoprotocatechuate dioxygenase is involved in the initial deoxygenation of 4-hydroxyphenylacetate to homoprotocatechuate (and eventually to fumarate). The monocyclic α subunit (Mono_DIOX) and bicyclic extradiol α subunit dioxygenases genes (EXDO_bi) and gentisate dioxygenase α subunit gene (Gen) from the cupin superfamily were also in the BONCAT+ metagenome (Fig. 4, Table S2), although at lower abundances across all shotgun metagenomes compared to the *lig*B dioxygenase (Fig. 5, Table S4). The *lig*B dioxygenase was found in the putative methanogenic BONCAT+ MAG that was most closely related to *Methanothrix* (discussed above) indicating possible activity of aerobic hydrocarbon degradation (*lig*B) in addition to anaerobic hydrocarbon degradation (via *pps*AB and *bss*D). Possible oxygen-dependent or -tolerant properties of this putative methanogenic population are discussed in detail in a separate study^48^. Further, the bicyclic dioxygenases and the gentisate dioxygenases (Gen) were in a BONCAT+ MAG taxonomically affiliated with Proteobacteria, while the monocyclic dioxygenases were detected in a different MAG within the β-Proteobacteria (Fig. 4, Table S2) indicating further aerobic degradation capabilities.

The anaerobic hydrocarbon degradation gene with the greatest abundance in the low-sulfate environments was *ppc*C and was identified in BONCAT+ MAG most closely related to *Geobacter* (Fig. 4, Fig. 5, Table S2 and Table S4). The high abundance of *ppc*C in both high- and low-sulfate environments indicates that anaerobic phenol degradation is likely crucial in the degradation of PRB coal. Phenylphosphate carboxylase genes are often overlooked for the more well documented fumarate addition genes^14,15,19,27,28^, but results presented herein suggest *ppc*C should be considered as a possible mechanisms for anaerobic hydrocarbon degradation. Phenol degradation via the activation of phenylphosphate followed by carboxylation has previously been shown to only occur when in the presence of phenol and CO_2_ and is believed to be a strictly anaerobic process^58^. However, previous research suggests *ppcC* can exist in an inactive oxidized form until it is reduced and activated under anaerobic conditions^61^. Therefore, under fluctuating oxygen conditions, phenylphosphate carboxylase may play an important role.

While all the fumarate addition genes of interest were identified in the environmental shotgun metagenomes (Fig. 5) and belonged to predominant populations related to the *Geobacter* (Bin 34), *Syntrophus aciditrophicus* (Bin 84), *Anaeromyxobacter* (Bin 37), *Luteitalea pratensis* (Bin 10), and *Desulfococcus* (Bin 368 and Bin 50) (Fig. 3 and Supplemental Data 2), only two of the 24 BONCAT+ MAGs displayed the presence of fumarate genes (related to *Methanothrix* and β-Proteobacteria) and of the seven fumarate genes investigated only three (*assD, bssD*, and *tutE*) were present (Fig. 4, Fig. 5, Table S2, Table S4). The functional significance and gene arrangement of these genes could be explored in future work. It is possible that the low-sulfate and methanogenic conditions were not able to support fumarate addition, due to the lack of available electron acceptors, which are necessary for fumarate addition and therefore needed to rely on other methods of carbon degradation.

Together, our results suggest that fumarate is an important intermediate substrate in hydrocarbon degradation for both high- and low-sulfate environments, but likely plays a more important role in high-sulfate PRB environments. Previous laboratory investigations have demonstrated an increase in *ass*A gene abundance during carbon degradation in sulfate-reducing environments, with no such increase in corresponding methanogenic cultures^15,19,21^. In contrast, other oil field surveys and hydrocarbon degradation experiments demonstrate that fumarate addition is an important pathway under methanogenic conditions and organisms with the *ass*A, such as *Anaerolineaceae*, have shown the ability to degrade *n*-alkanes to produce acetate, potentially creating a syntrophic relationship with the acetoclastic methanogen, *Methanothrix*^62,23,26^. Our observations suggest that in low-sulfate coal seams, prerequisite methanogenic substrates may result from both anaerobic and aerobic (homoprotocatechuate dioxygenase) hydrocarbon degradation as they are detected in higher gene abundancess across all coal-dependent and active populations (Fig. 4, Fig. 5, Table S2, and Table S4).

## Conclusion

The microbial environment of the terrestrial subsurface is one of the most complex and poorly understood microbiomes on Earth^63^. Due to diversity and high functional capacity, there have been many calls to better understand the diversity^64^ and the activity of *in situ* microorganisms and the potential implications on the cycling of old and new carbon between the lithosphere and atmosphere^65^. There has been much debate on possible pathways actively involved in the subsurface carbon cycle, specifically, the degradation of complex hydrocarbons to methanogenic substrates under varying conditions (*e*.*g*. sulfate). The subsurface environment constitutes a challenging environment to access and prior to this study, conclusions had been based primarily on laboratory benchtop enrichments, geochemical predictions, or shotgun metagenomic models^2,5,24^. The methods used in this work, accessing the subsurface and sampling of environmentally relevant *in situ* communities using the SES and the integration of metagenomes from active communities derived from BONCAT-FACS, expands both the gene and genome catalog of the subsurface environment as well as the activity of key carbon cycle genes indicative of hydrocarbon degradation. The active groups were putatively involved in both aerobic and anaerobic hydrocarbon degradation and biosurfactant production. The presented results demonstrated that active populations may have the ability to produce diverse biosurfactants and could play an important role in coal-dependent methanogenesis in low-sulfate coal seams. While this study proposes biosurfactants are important in low-sulfate coal seams, more work needs to be performed to determine the role and environmental triggers for biosurfactants from different coal ranks and sulfate concentrations.

Hydrocarbon degradation in coal seam environments is often assumed to be strictly anaerobic, although with advancements in metagenomics more research is beginning to suggest mixed aerobic and anaerobic hydrocarbon degradation^24,66^. Our results indicate that aerobic hydrocarbon degradation genes are associated with active populations in coal seam environments that span redox zones^8^ and are more prevalent in low-sulfate coal seams. Previously, enzymes belonging to the LigB superfamily had not been readily detected due to conventional screening techniques. In the advent of metagenomics, these enzymes have been broadly identified in the environment with an enrichment in hydrocarbon rich samples, suggesting enhanced importance in hydrocarbon degradation^67,68^. The BONCAT+ metagenomes were predominated by both aerobic and anaerobic degradation genes that included *ppc*C, *pps*A, *ebd*B, *cmd*B, *ass*D, *lig*B, monocyclic and bicyclic dioxygenases. Therefore, based on the active MAGs that contain aerobic genes, it is possible microaerophilic conditions could promote aerobic hydrocarbon degradation via dioxygenases such as the homoprotocatechuate dioxygenases from the LigB family. The source of oxygen in these environments is unknown, and likely depends on groundwater flow/recharge, surface water intrusion, water level changes, coal seam depth, and associated microbial activities^24,47^. Further work should determine the extent of oxia, potential oxygen sources, and potential segregation of hydrocarbon degradation and methanogenesis along heterogeneous critical zone transitions in the shallow subsurface.

## Methods

### Site Description and Sample Collection

The USGS Birney sampling site, located in the PRB of southeastern Montana (45.435606, -106.393309), has access to four subbituminous coal seams at different depths and within a 15 meter lateral distance of each other^8^. Metagenomic samples were collected from a high-sulfate (23.78 – 26.04 mM) coal seam (N-H) at 65 meter depth and two low-sulfate (0.01 – 0.38 mM) coal seams (FG-L and T-L) located at 117 meter and 161 meter depths. The low-sulfate coal seams have measurable methane (33 – 67 mg/L) while the high-sulfate coal seams have low methane (<0.15 mg/L) as previously measured^8^. The hydrogeochemistry of these coal seams appear to remain relatively stable with sulfate and methane measurements fluctuating very little between 2011 and 2014^8^ (Table S5). Coal associated microbial assemblages were collected with a diffusive microbial sampler (DMS) as previously described^8^ or with a next-generation subsurface environmental sampler (SES; Patent# US10704993B2) following previous field protocols. The DMSs were incubated down-well for three months and retrieved on June 12^th^, 2017. Once retrieved, the slurry (coal and groundwater) from the DMSs were aseptically removed and stored on dry ice in sterile falcon tubes until brought back to the lab, where they were stored at -80°C until DNA extractions were performed.

### DNA Extraction, Concentration and Sequencing for Shotgun Metagenomics

DNA was extracted from slurry in the DMS as previously described in Schweitzer et al.^5^ using a FastDNA Spin Kit for Soil (MP Biomedical). DNA was purified using One Step PCR Clean Up (Zymo Research). The DNA was quantified using Qubit dsDNA HS Assay Kit (Invitrogen) before being sent to the MBL Keck sequencing facility (The University of Chicago, IL). The genomic DNA was concentrated using a SpeedVac (Thermo Scientific) and quantified using Picogreen (Invitrogen). DNA concentrations ranged between 138.6ng (FG-L) and 1,450ng (T-L). Using a Covaris ultrasonicator, DNA was sheared to ∼400 bp and libraries were constructed with NuGEN Ovation^®^ Ultralow Library protocol (Tecan Genomics, CA). The amplified libraries were visualized on a Caliper HiSense Bioanalyzer and pooled to equimolar concentrations. The DNA was further size selected using a Sage PippinPrep 2% cassette. The pooled library was quantified using Kapa Biosystems qPCR library quantification before being sequenced in a 2×150 paired-end sequencing run using an Illumina NextSeq. The sequences were demultiplexed and the adapter sequences were trimmed using bcl2fastq conversion software. The sequences were archived with SRA (SRP292435) under the BioProject ID PRJNA678021.

### BONCAT Incubations, Fluorescent Labeling, Cell Sorting and Metagenomic Sequencing

The following methods are also presented in our companion study, which examines four high-quality BONCAT+ MAGs for their involvement in acetate production and acetoclastic methanogenesis^48^. Subsurface environmental samplers (SES, Patent # US10704993B2) that contained UV-sterile, crushed coal (size range 0.85 – 2.00 mm) were deployed at the screened interval in a methane-producing coal seam (FG-L; September 2017) at the USGS Birney Field Site^8^ and, after approximately nine months of down-well incubation, were retrieved. In contrast to DMSs used for shotgun sequencing, the use of the SES allowed for samples to be kept under anaerobic conditions and at formation pressures, an essential prerequisite for conducting representative BONCAT incubations. Upon retrieval, 10 mL of SES slurry was sampled via sealed ports (Swagelok) and were anaerobically transferred into gassed-out, sterile Balch tubes. Triplicate tubes were prepared for incubation with 250 μM L-homopropargylglycine (HPG, Click Chemistry Tools, Scottsdale, AZ, USA) prepared in sterile, degassed water (DEPC diethyl pyrocarbonate treated, pH 7). Control samples were prepared the same as other samples with the exception that HPG was not added (HPG negative control). Samples were incubated in the dark at 20 °C for 24 hrs. At the end of the incubation period cells were removed from coal by removing 1 mL of slurry and adding 5mls Tween^®^ 20 dissolved in PBS (1X PBS) to a final concentration of 0.02% (Sigma-Aldrich). Samples were vortexed at maximum speed for 5 min followed by centrifugation at 500 × g for 5 min (Couradeau et al., 2019)^38^. The supernatant (containing the detached cells) was immediately cryopreserved at −20 °C in sterile 55% glycerol TE (11X) solution.

Translationally active cells were identified through a click reaction that added a fluorescent dye to HPG molecules that had been incorporated into newly synthesized protein. This BONCAT click reaction consisted of 5mM sodium ascorbate, 5mM aminoguanidine HCl, 500uM THPTA, 100uM CuSO_4_, and 5uM FAM picolyl azide in 1X phosphate buffered saline. Incubation time was 30 minutes, followed by three washes in 20ml of 1X PBS for 5 minutes each. Cells were recovered from the filter by vortexing in 0.02% Tween for 5 minutes, and then stained using 0.5uM SYTO™59 (ThermoFisher Scientific, Invitrogen, Eugene OR, USA) DNA stain (Couradeau et al., 2019)^38^.

Cells were sorted using a BD-InfluxTM (BD Biosciences, San Jose, CA, USA) configured to capture total cells labeled with SYTO™59 DNA stain and excited with a 640nm red laser and BONCAT-positive cells labeled with FAM picolyl azide dye and excited with a 488nm blue laser. BONCAT-positive cells were a subset to total SYTO™59 stained cells, and were differentiated from the BONCAT-negative based on their FAM fluorescence (530/40BP) to background fluorescence from identical cells inclubated without HPG, *i*.*e*., HPG negative control, that also underwent the same click reaction to add a FAM label. The two populations were sorted. The first population contained all DNA+ cells (BONCAT-total), and the second population contained only those cells which were also BONCAT+ in comparison to the control. For each sample, we sorted 4 wells containing 5,000 cells each, and 20 wells containing 300 cells each into 384 wells plates. Plates were frozen at −80 °C until further processing.

Cells from samples containing 5,000 sorted cells were pelleted prior to sequencing via centrifugation at 6,000 x g for 1 hour at 10 °C. The supernatant was discarded by a brief inverted spin at <60 x g so as not to interfere with subsequent whole genome amplification reaction chemistry. Plates containing only 300 sorted cells were not pelleted. Sorted cells were lysed and amplified using 5 µl WGAX reactions following the optimized conditions described in Stepanauskas et al. (2017)^69^. Briefly, cells were lysed in 650 nl lysis buffer for 10 minutes at room temperature. The lysis buffer consisted of 300 nl TE + 350 nl of 400 mM KOH, 10 mM EDTA, 100 mM DTT. Lysis reactions were neutralized by the addition of 350 nl of 315 mM HCl in Tris-HCl. Amplification reactions were brought to 5 µL with final concentrations of 1X EquiPhi29 reaction buffer (Thermo), 0.2U/µl EquiPhi29 polymerase (Thermo), 0.4mM dNTPs, 50µM random heptamers, 10 mM DTT, and 0.5 µM SYTO13. Plates were incubated at 45 °C for 13 hours.

Libraries for BONCAT+ and BONCAT-total metagenomic sequencing were created using the Nextera XT v2 kit (Illumina) with 12 rounds of PCR amplification. All volumes and inputs to Nextera reactions were reduced 10-fold from the manufacturer’s recommendations. Libraries were sequenced 2X150bp mode on Illumina’s Nextseq platform. The sequences were archived under JGI GOLD Study ID Gs0141001.

### Metagenomic Analysis with Anvi’o

NextSeq-generated fastq files were filtered at a minimum quality read length of 0.75 using Anvi’o for Illumina (http://merenlab.org). The filtered reads were co-assembled using MEGAHIT v1.1.2 (The University of Hong Kong and L3 Bioinformatics Limited). All shotgun, BONCAT+, and BONCAT-Total replicates were co-assembled and kept separate for subsequent comparisons. Alignments were accomplished using bowtie2 v2.3.6 and samtools v1.9. All bam files were merged into individual profile databases for each sample prior to binning using Anvi’o. Bins were assembled using tetranucleotide frequency into 376 genomic clusters for the shotgun metagenomics samples and 20 bins for the BONCAT+ amplified metagenome and 38 bins for the BONCAT-Total amplified metagenome. To determine completeness, scanning of ribosomal single copy genes for bacteria (Campbell et al. 2013^70^) and archaea (Rinke et al. 2013^71^) in each cluster was performed in Anvi’o. To identify open reading frames in contigs, Prodigal was used within Anvi’o. Functions were assigned in Anvi’o using NCBI’s Clusters of Orthologous Genes (COGs) and a Hidden Markov Model hit database was generated with Anvi’o. Coverage estimates, GC content, and N50 were calculated. The phylogenetic tree was created using MUSCLE (drive5.com/muscle) for multiple sequence alignment. Anvi’o Interactive Server (anvi-server.org) was used to create phylogenomic evaluation of genomic parameters for each MAG. All shotgun metagenome high quality (>80% completion), BONCAT+ and BONCAT-Total MAGs are archived under JGI GOLD (available upon publication).

### Metagenomic Analysis with Curated Database

Taxonomic identity was determined using NCBI BLASTn of all contigs for each bin. All blasted contigs with the best e-value and percent identity were then compared to select the most common ancestor to represent the most likely taxonomic identification for all the contigs for MAGs > 20X coverage. In order for a comprehensive assessment of up to date microbial hydrocarbon degradation strategies genes were analyzed using curated databases via direct protein sequence comparison. Curated databases for hydrocarbon degradation genes included aerobic (AromaDeg)^60^ and anaerobic (AnHyDeg)^20^ protein databases. Other functional gene databases (*e*.*g*., biosurfactant genes, *dsr*A, and *mcr*A) were generated from protein sequences by extracting HMM files from the Pfam database (EMBL-EBI) and from McKay et al. (2017).^13^ These protein databases were BLAST-searched against each protein sequence with an e-value cutoff of 1E-10 and above 40% identity. All duplicate gene hits were removed with the highest e-value hit being kept. All protein gene sequence coverages were normalized to give gene abundance by using Reads (Coverage) per kilobase of transcript per million mapped reads (RPKM).

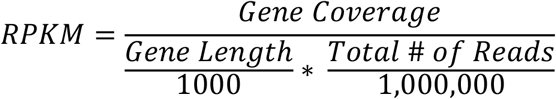

Genes were selected from the complete list of all genes (Table S4) and classified as genes of interest for further analyses. The genes of interest were selected first by ensuring there were representatives from each of the four databases tested (AnHyDeg, AromaDeg, Biosurfactant, and Redox) and that only one subunit representative from the same gene family was further analyzed. To determine which of the gene subuntit representatives were selected as a gene of interest, at least one of the following criteria had to be met: the gene had to either be identified in the BONCAT+ sample or contained the highest gene abundance in one of the samples (FG-L, T-L, or N-H).

## Data Availability Statement

All data is publicly available and archived under JGI GOLD Study ID Gs0141001.

## Acknowledgements

Metagenome sequencing was made possible by the Deep Carbon Observatory’s Census of Deep Life and was performed at the Marine Biological Laboratory (Woods Hole, MA, United States) and Joint Genome Institute CSP 503725. We are grateful for the assistance of Mitch Sogin, Joseph Vineis, Amelia Brumbaugh, and Hilary Morrison at MBL. The work conducted by the U.S. Department of Energy Joint Genome Institute, a DOE Office of Science User Facility, is supported under contract No. DE-AC02-05CH11231. Any use of trade, firm, or product name is for descriptive purposes only and does not imply endorsement by the U.S. Government. The US Geological Survey (E.P.B) supported field work and sampling. The work was further supported by NSF 1736255 (L.J.M., R.G., and M.W.F.). The authors would also like to acknowledge Dr. Katie Davis and George Platt for their help in sample collection in the Powder River Basin.

## Author Contributions

H.D. Schweitzer, H.J. Smith are coauthors and contributed equally to the manuscript. H.D. Schweitzer, H.J. Smith, E.P. Barnhart, A.B. Cunningham, and M.W. Fields designed the study. H.D. Schweitzer, H. Smith, and E.P. Barnhart wrote the paper. H.D. Schweitzer, H.J. Smith, and E.P. Barnhart collected samples from the field. H.D. Schweitzer, H.J. Smith, and E.P. Barnhart performed experiments. R.R. Malmstrom and D. Goudeau performed cell sorting and sequencing. H.D. Schweitzer, and H.J. Smith performed analysis. All authors critically reviewed the manuscript. M.W. Fields supervised research.

