## Supplemental Legends and Info for "Subsurface Hydrocarbon Degradation Strategies in Low- and High-Sulfate Coal Seam Communities Identified with Activity-Based Metagenomics"

**Affiliations:**

[2]Department of Microbiology & Cell Biology, Montana State University, Bozeman, MT 59717, USA

[†]Now at UiT - The Arctic University of Norway, 9019 Tromsø, Norway

H.D. Schweitzer, Post Doctoral Researcher

UiT - The Arctic University of Norway

The Norweigian College of Fishery Science

Muninbakken 21

9019 Tromsø, Norway

M.W. Fields, Professor

Montana State University

Center for Biofilm Engineering

366 EPS Building

Bozeman, MT 59717, USA

**Supplemental Information Contents:**

Supplemental Figures S1-S3

Supplemental Tables 1-5 and Supplemental Data 1-3

**Supplemental Figures:**

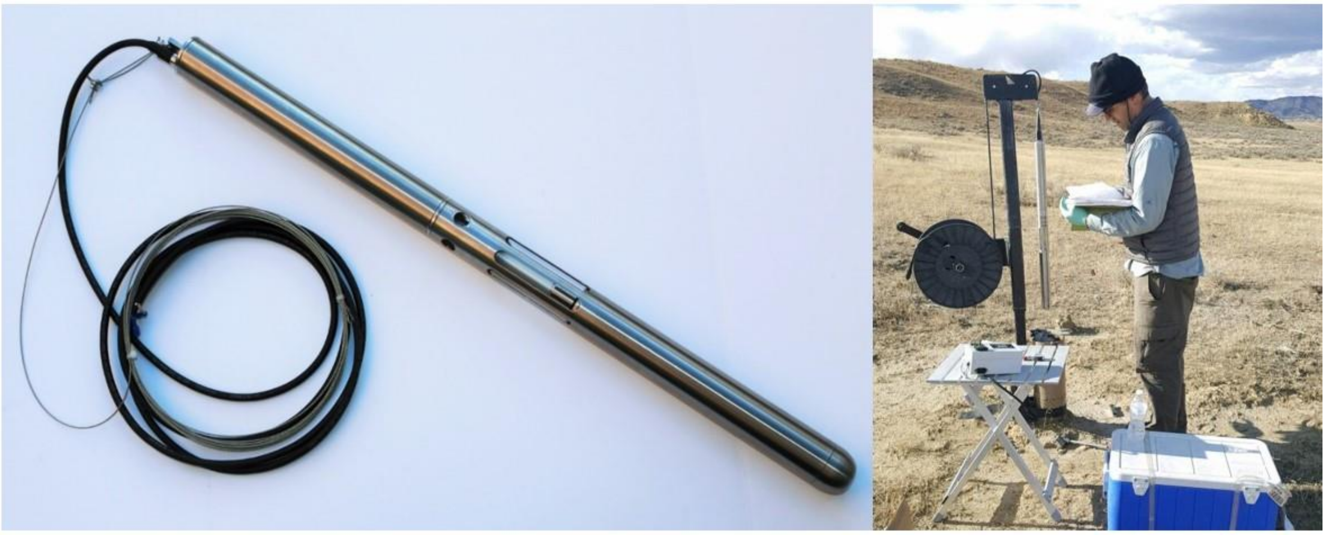

Supplemental Figure S1.) Photographs of the Subsurface Environmental Sampler (SES) used to obtain coal associated communities.

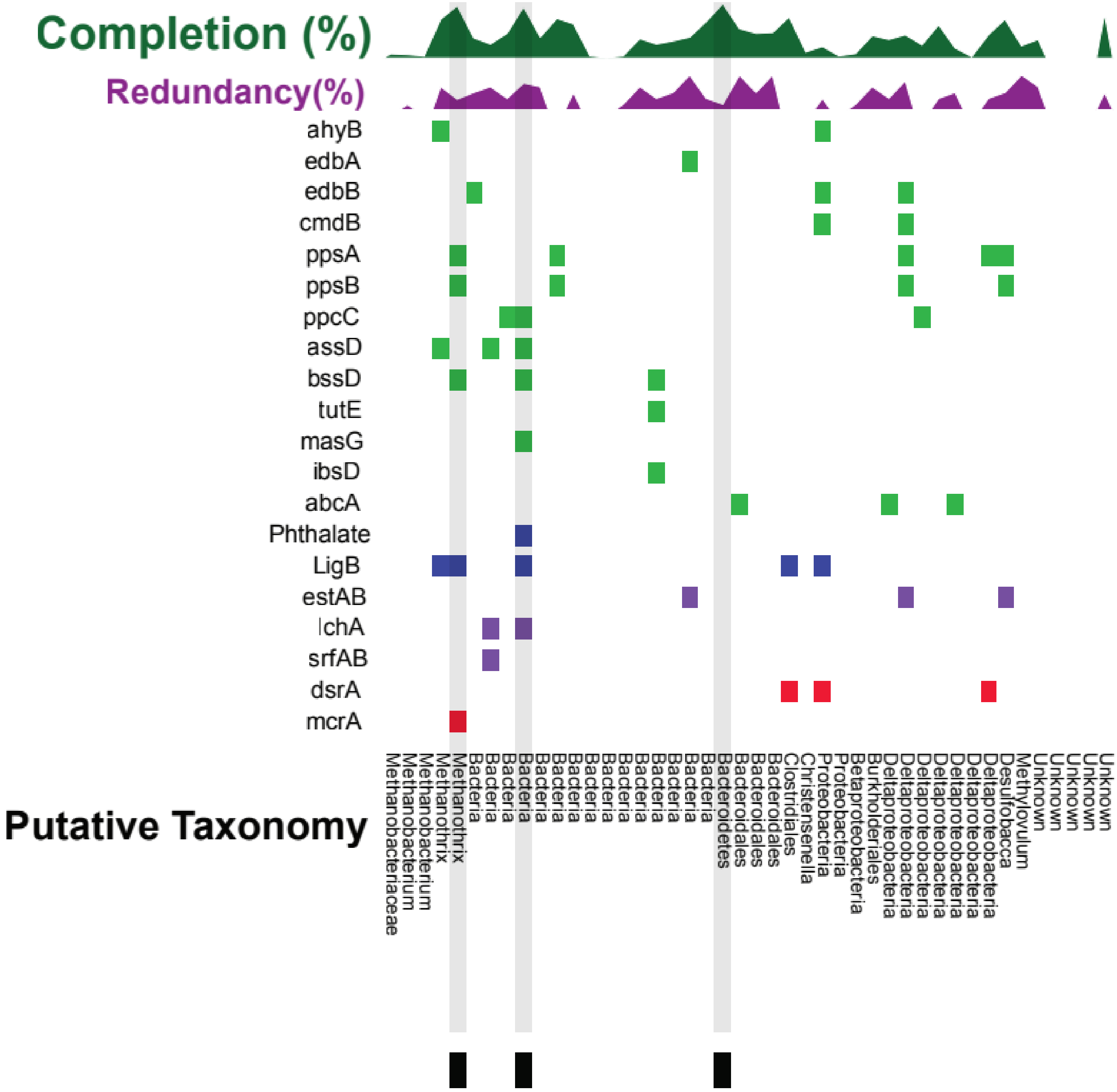


Supplemental Figure S2.) Evaluation of SYTO-Total metagenome assembled genomes (MAGs) and the presence or absence of anaerobic (green) and aerobic hydrocarbon degrading (blue), biosurfactant (purple) and sulfate reduction/methanogenesis (red) genes for each corresponding MAG. The AnHyDeg and AromaDeg databases were used to determine the different hydrocarbon degradation genes used for comparison. The percent completion (green line graph out of 100%) and redundancy (purple line graph out of 10%) were compared for each MAG. MAGs with putative taxonomy Methanothrix, Bacteria and Bacteriodetes were all greater than 92% complete and contain an average nucleotide identity that is >99% similar to their matching BONCAT MAGs which is represented by the symbol (■).

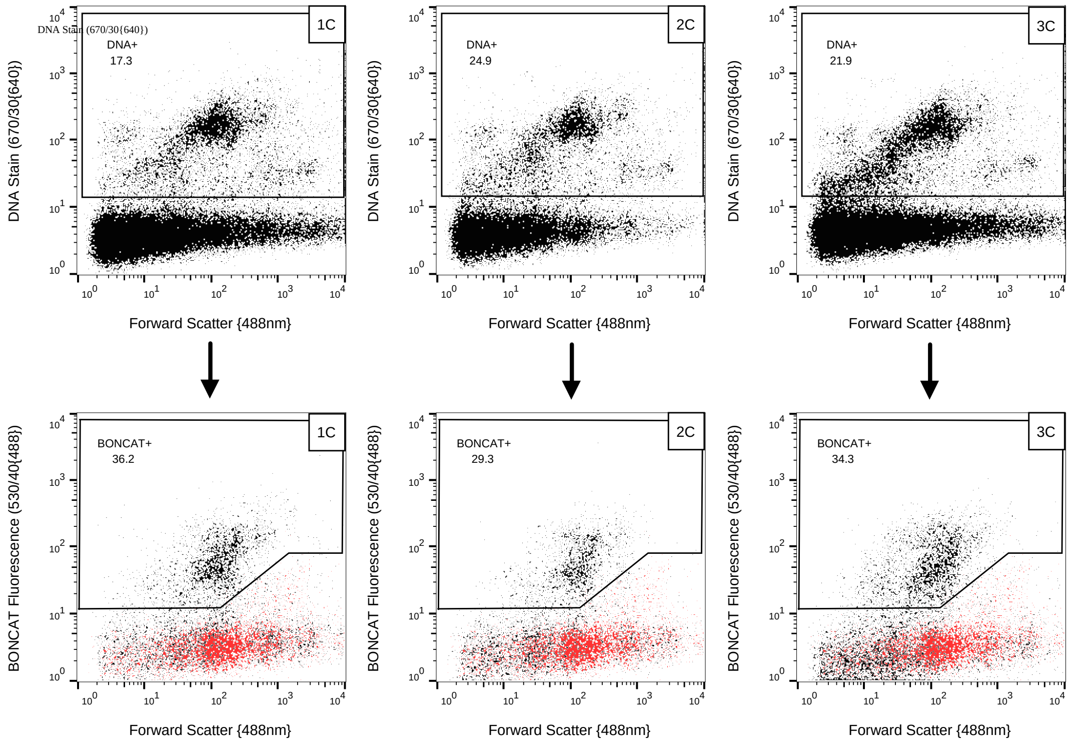


Supplemental Figure S3.) Gate drawing accomplished in two steps for the biological replicates of FG-11 BONCAT positive samples. The top panel 1A, 2A, and 3A is gated to separate total cells from background particles and gates indicated in black are based on a DNA stain (Syto59, ex: 640nm em: 655-685nm). Following the initial gating, SYTO+ cells were analyzed further for BONCAT fluorescence with the FAM (Picolyl dye (Ex: 488nm/Em: 530nm) (bottom panel). Gate for 1B, 2B, and 3B. Gating of the BONCAT+ populations was achieved by comparing each replicate to both HPG negative and water controls (indicated in red overlaid in each graph). The number in the top left of each of the bottom panel graphs indicates the percentage of BONCAT+ cells in comparison to total cells (SYTO+) for each replicate.

**Supplemental Tables/Data:**

***Uploaded as a data file with table explanations provided below**

Supplemental Table 1.) List of environmental metagenome assembled genomes (MAGs) from high sulfate and low sulfate coal seams from the Powder River Basin and the corresponding sequencing and analysis parameters.

Supplemental Table 2.) List of BONCAT+ metagenome assembled genomes (MAGs) and the corresponding sequencing and analysis parameters. The corresponding genes present in each MAG are listed under their corresponding database.

Supplemental Table 3.) List of SYTO-Total metagenome assembled genomes (MAGs) and there corresponding sequencing and analysis parameters. The corresponding genes that are present in each MAG are listed under their corresponding database.

Supplemental Table 4.) Complete list of the four databases and all the genes compared in this study. The function of each gene and the coverage of each gene in each sample is listed and the presence of gene in the BONCAT+ sample is indicated. The genes further analyzed are marked as genes of interest.

Supplemental Table 5.) Formation water and dissolved gas chemical and isotopic composition. Proximate/ultimate analysis percentages and heating values of the coal. All data reported in this table was modified and described in more depth in Barnhart et al. 2016.

Supplemental Data 1.) List of functional genes of all samples from the Kyoto Encyclopedia of Genes and Genomes (KEGG) Gene database.

Supplemental Data 2.) The gene abundances (calculated with RPKM) of individual genes of interest involved in hydrocarbon degradation for the shotgun environmental MAGs from the high-sulfate coal seam (N-H) and two low-sulfate coal seams (T-L and FG-L). All MAGs that were below 80% completion were grouped together as <80% Completion.

Supplemental Data 3.) Average Nucleotide Identity comparison of all shotgun, BONCAT+ and SYTO-Total metagenomes.
