## Supplementary figures and images for "Subsurface Hydrocarbon Degradation Strategies in Low- and High-Sulfate Coal Seam Communities Identified with Activity-Based Metagenomics"

### Supplemental Figure 1

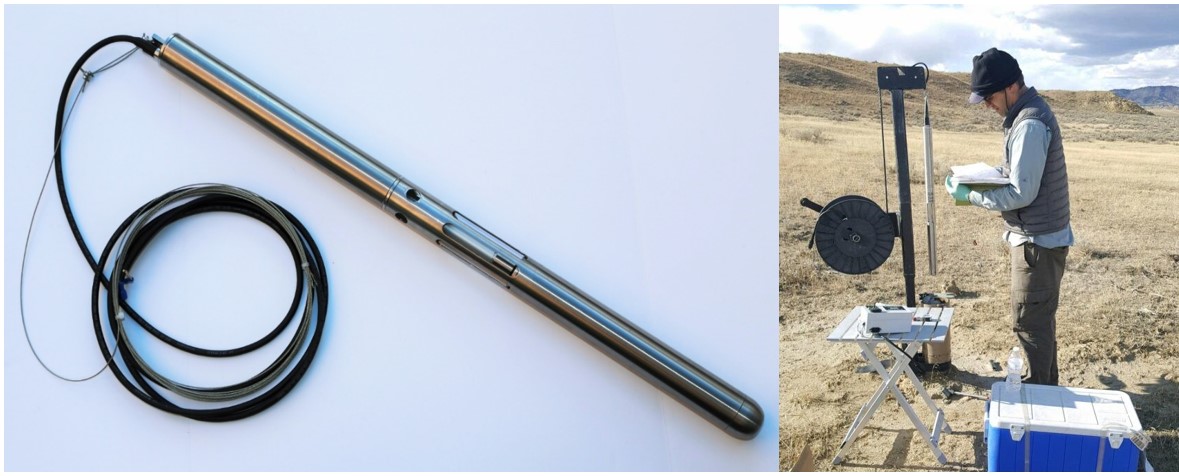

### Supplemental Figure 2

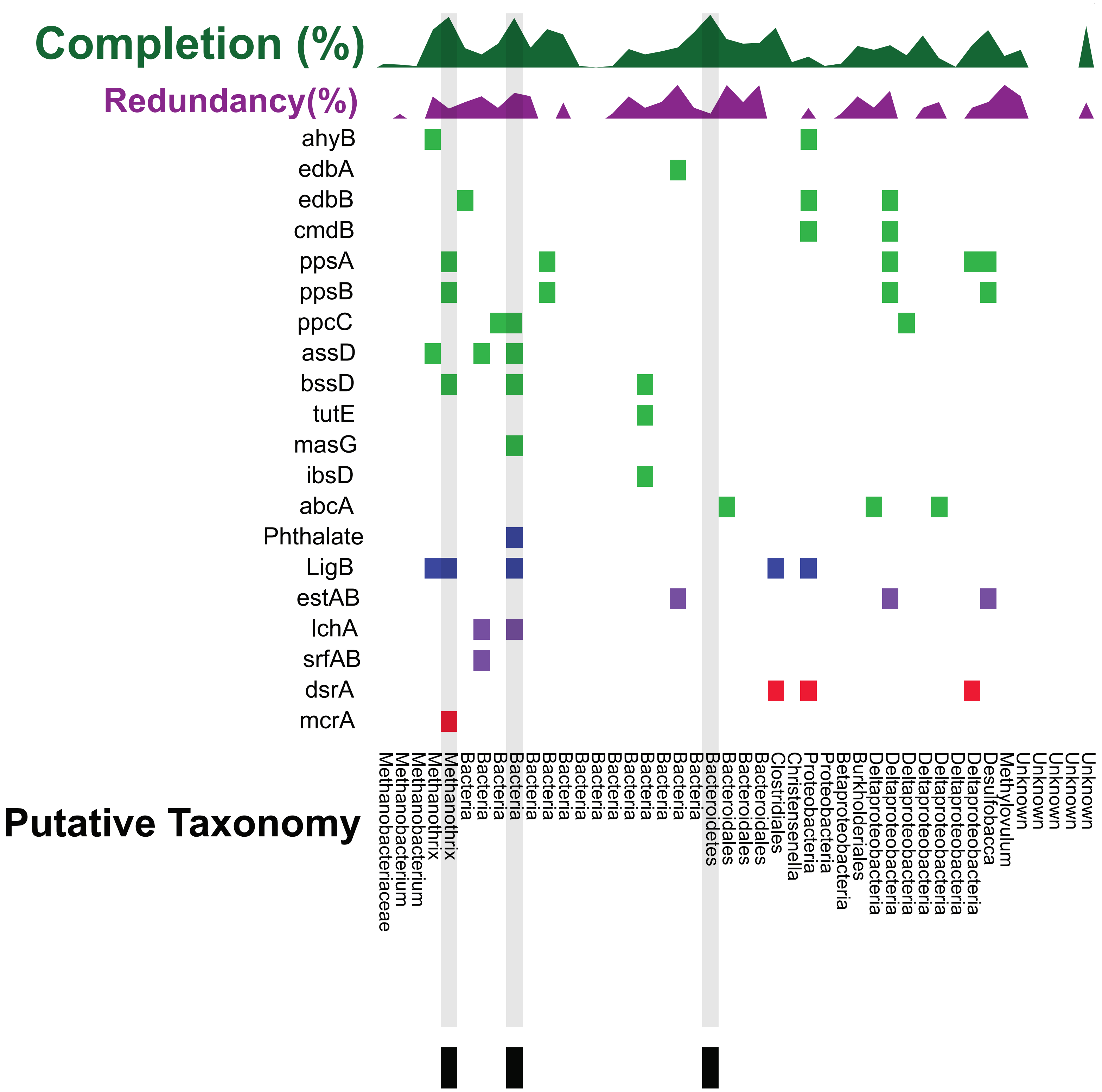

### Supplemental Figure 3

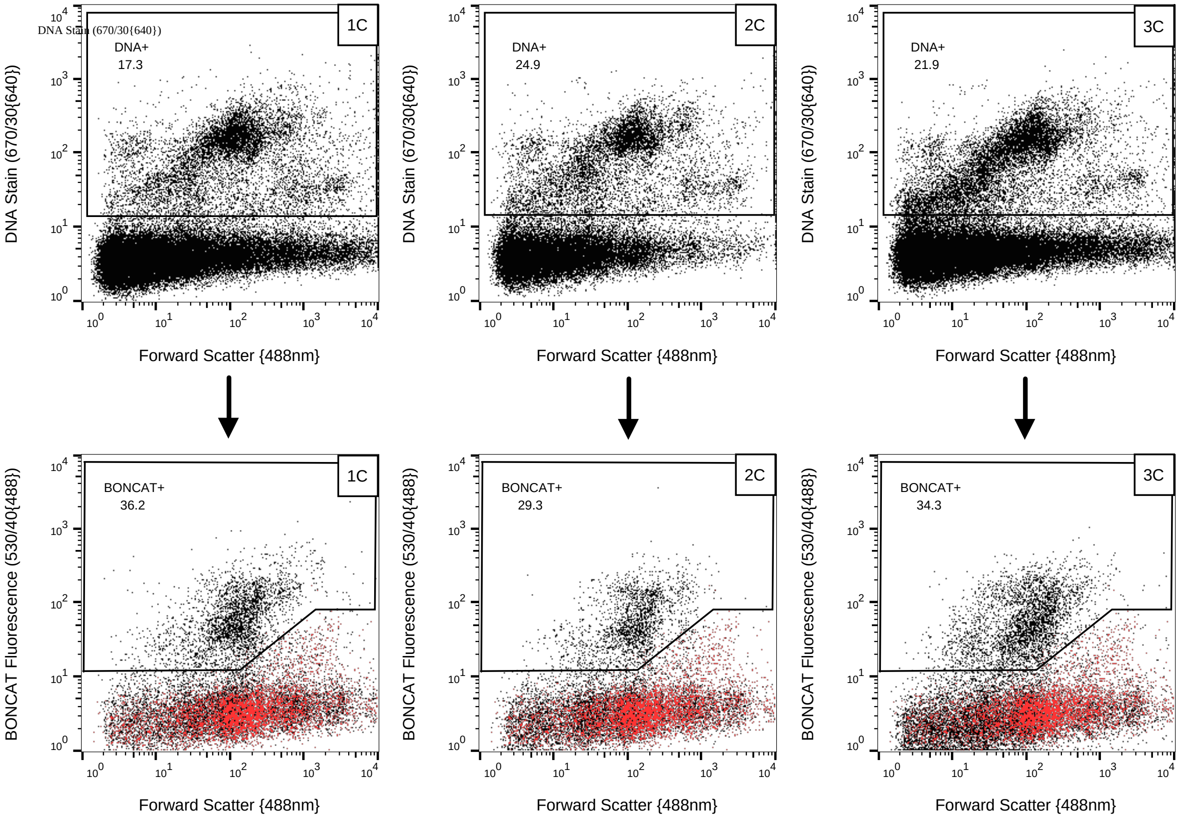
